# Characterising Complex Enzyme Reaction Data

**DOI:** 10.1101/028142

**Authors:** Handan Melike Dönertaş, Sergio Martínez Cuesta, Syed Asad Rahman, Janet M. Thornton

## Abstract

The relationship between enzyme-catalysed reactions and the Enzyme Commission (EC) number, the widely accepted classification scheme used to characterise enzyme activity, is complex and with the rapid increase in our knowledge of the reactions catalysed by enzymes needs revisiting. We present a manual and computational analysis to investigate this complexity and found that almost one-third of all known EC numbers are linked to more than one reaction in the secondary reaction databases (e.g. KEGG). Although this complexity is often resolved by defining generic, alternative and partial reactions, we have also found individual EC numbers with more than one reaction catalysing different types of bond changes. This analysis adds a new dimension to our understanding of enzyme function and might be useful for the accurate annotation of the function of enzymes and to study the changes in enzyme function during evolution.

## Introduction

Enzymes are life’s catalysts that accelerate biochemical reactions up to the rates at which biological processes take place in living organisms. They play a central role in biology and have been thoroughly studied over the years. Since the 1960s, the Nomenclature Committee of the International Union of Biochemistry and Molecular Biology (NC-IUBMB) has systematically encapsulated the functional information of enzymes into EC numbers. Considered in some cases as an enzyme nomenclature and classification system, the EC is one way to annotate enzymes, by a classification of the representative reaction they catalyse, based on multiple aspects of the overall chemistry such as the chemical bonds that are broken or formed, cofactors being used and the nature of the substrates undergoing transformation. Introduced into the widely used Gene Ontology (GO) system for the functional annotation of genes, the EC is the global standard representation of molecular function for enzymes and relates biological information such as genes, sequence and structure with chemistry data in resources like UniprotKB [1].

The EC classification as defined by IUBMB is a primary resource for information about enzyme function. Other databases such as KEGG [2] and BRENDA [3] are based around the IUBMB definitions, however in order to handle the flood of data, they associate additional reactions to EC numbers at their discretion, which sometimes causes problems. Nevertheless, the EC has proved to be very powerful. It is manually curated and maintained by expert enzymologists, who use a controlled vocabulary and well-defined relationships in describing enzyme function [4] to convey the way biochemists think about reactions [5]. It facilitates predefined comparisons between enzymes reactions and newly discovered reactions are easily allocated in the different levels of its hierarchical classification. However, because of the diversity of chemical criteria used at different levels, the classification is not coherent between EC classes [6-8]. For instance, lyases (EC 4) are divided in subclasses depending on the type of chemical bond that is broken whereas isomerases (EC 5) are divided based on the type of isomerisation. In addition, the EC classification is based on the overall catalysed reaction, which means that mechanistic steps and reaction intermediates are not considered. As a result, enzymes carrying out the same overall reaction are generally assigned to the same EC number, even when they perform catalysis using different cofactors and mechanisms [9]. For example, three structurally distinct non-homologous chloride peroxidases, which are deemed to have emerged from independent evolutionary events [10,11], catalyse the chlorination of alkanes using three different mechanisms and cofactors. However they are all associated to the same EC number (EC 1.11.1.10). First, vanadate is a prosthetic group in an acid-base mechanism [12] [13]. Second, heme is also a prosthetic group in a radical mechanism [14]. Third, a Ser-His-Asp catalytic triad and an organic acid cofactor are involved in an acid-base mechanism [15]. On the other hand, enzymes catalysing the same overall reaction using the same mechanism with slightly different cofactors are sometimes assigned different EC numbers. For instance, EC 1.1.1.32 and 1.1.1.33 represent two mevaldate reductases, both catalyse the conversion of (R)-mevalonate to mevaldate but respectively use NAD^+^ and NADP^+^ as a cofactor [16].

Although reliable and rigorous, the manual process of naming each new enzyme and classifying novel enzyme reactions is laborious and requires expert knowledge, therefore automatic approaches may help to accelerate this procedure and to guide the navigation between related enzyme reactions. Similarly, the IUBMB has also considered the current EC classification system to be a relic of the original attempts to develop a chemically sensible hierarchical classification. Ideas and methodologies envisioning a new system in which enzymes are assigned meaningless database identifiers have already been proposed [17] and automatic tools to search and compare enzyme reactions are useful to navigate through “enzyme reaction space” and may help to improve future versions of the classification [18].

There are biological aspects of enzyme function that are hard to capture in a hierarchical classification system [19]. First, enzymes can be promiscuous and catalyse more than one biochemical reaction [20]. Second, homologous enzymes annotated with the same EC number can manifest different levels of substrate specificity [21] (also known as substrate promiscuity or ambiguity). For instance, UDP-glucose 4-epimerases (EC 5.1.3.2) display different substrate specificities depending on the taxonomic lineage. Bacterial epimerases only act upon UDP-glucose whereas the eukaryotic relatives additionally catalyse the transformation of UDP-N-acetylglucosamine [22]. Even though this limitation has partially been addressed by introducing specificity information in the “Comments” section of several EC entries [23], there is still a need to represent this phenomenon in a more computer-friendly format in order to obtain accurate comparisons between EC numbers. Third, the inclusion of enzyme sequence and structural information would add biological insight to the EC assignment process [21]. This is particularly severe when classifying enzyme functions that involve polymeric biomolecules like sugars, proteins or DNA. For instance, proteolytic and carbohydrate-active enzymes exhibit broad substrate specificity and have been alternatively classified using sequence and structure analyses in the MEROPS [24] and CAZy [25] resources. Fourth, more than 30% of all EC numbers are orphans, where no enzyme information is known at all [26]. This represents a challenge for the accurate interpretation of enzyme function in high-throughput sequencing initiatives.

Evidence suggests that the correspondences between enzymes, EC numbers and reactions are not simple [19,27]. The relationship between enzyme and EC number is complex and rarely one-to-one [10]. Some enzymes are annotated with multiple EC numbers (multifunctional) [5] whereas some EC numbers are associated with many unrelated enzymes [11]. For example, several studies have deliberatively excluded multifunctional enzymes in order to avoid complexities [28,29]. The relationship between EC number and reaction is not straightforward either. Although the IUBMB definitions are the standard, there are striking differences in the way reactions are represented using the EC classification in several databases. The majority of biologists use the KEGG database in their work to look at reactions because it provides easy access to chemical equations and molecular structures for academic users and it is complete in comparison with other databases. Although various studies exclude reactions associated with more than one EC number [30,31], some approaches aiming to predict reactivity in metabolites have successfully handled reactions associated with more than one EC number [32]. To some extent, KEGG circumvents the need for using EC numbers to link enzymes and biochemical reactions by directly connecting reactions to groups of orthologous enzymatic genes [33]. This association might considerably simplify the process of linking chemical and genomic information in the future.

This study examines the complexity in the relationship between EC number and reaction in the KEGG database. Although some reviews mentioned aspects of this connection [26,34], to the authors’ best knowledge, studies addressing its complexity in a systematic manner are lacking. We first explored this relationship for a chemically diverse class of enzymes catalysing geometrical and structural rearrangements between isomers, the isomerases. Although this class accounts for only 5.2% of all EC numbers, their diverse chemistry and the similarity of some subclasses to EC primary classes [35], makes the isomerases a class which is representative of the overall chemistry of the EC classification. The knowledge derived from the manual analysis was used to develop an automatic approach to gain an overview of reaction diversity across the EC.

## Methods

### Overview

There are 5385 four-digit EC numbers in the 9th April 2014 release of the NC-IUBMB list, 4237 of them (79%) are associated with 6494 unique reactions bearing structural information in the 70.0+ release of KEGG database [2], accessed using the KEGG website and Advanced Programming Interface (API) [36]. The remaining 21% lack structural data. Although most EC numbers are linked to one reaction, almost a third are associated with more than one (Fig. 1a). Comparatively, oxidoreductases (EC 1) exhibit the highest fraction of multiple reactions whereas isomerases (EC 5) the lowest (Fig. 1b). Similarly, some unusual cases were identified where individual EC numbers are linked to over 20 reactions, with one extreme outlier, classified as an unspecific monooxygenase (EC 1.14.14.1) with up to 66 reactions (Fig. 1c). In isomerases, the total number of EC numbers in the database is 245, for which 222 are associated with 298 biochemical reactions and 23 are not linked to any reaction. Among the EC numbers linked to isomerase reactions, 42 are associated with more than one reaction.

**Fig. 1.**
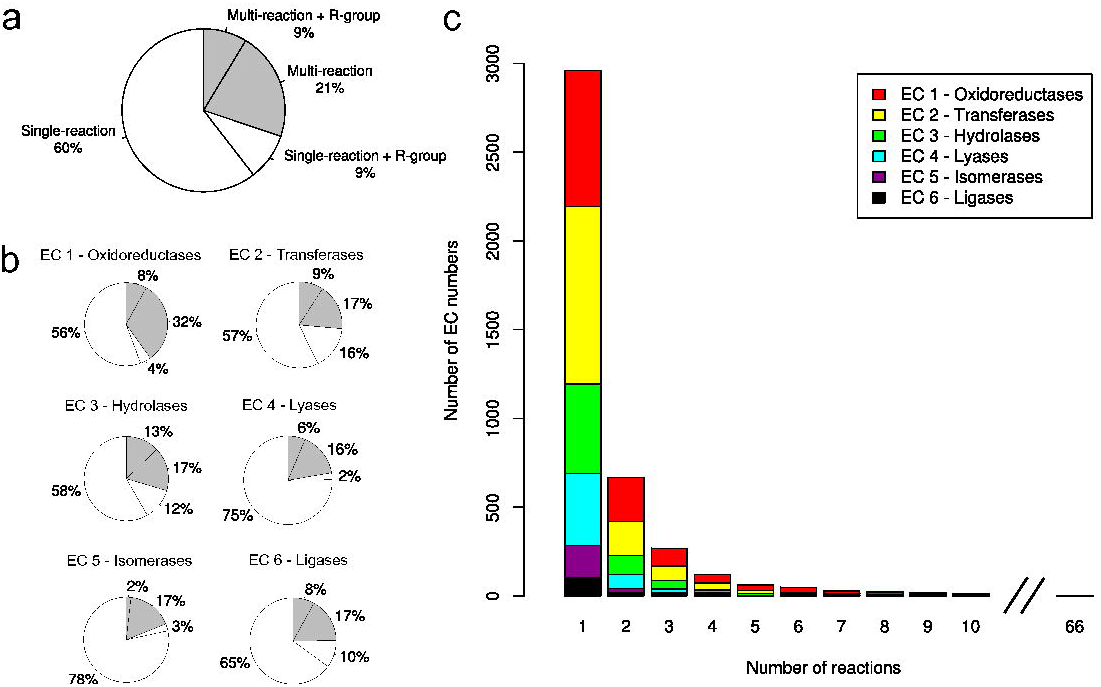
Survey of EC numbers associated with more than one enzyme reaction. (a) Overall distribution. White and grey slices indicate single and multi-reaction EC numbers, respectively. “R-group” represents EC numbers containing a Markush label in at least one reaction (see *Generic* reactions in main text) (b) Distribution by EC class (c) Distribution of EC numbers according to the number of reactions.

### Automatic analysis – extending diversity groups found in isomerases to the EC classification

The automatic extraction of chemical attributes from biochemical reactions such as bond changes is necessary to compare enzymes based on the chemistry of their catalysed reactions. In order to calculate chemical attributes we used EC-BLAST, a recently-developed algorithm to obtain accurate atom-atom mapping, extract bond changes and perform similarity searches between enzyme reactions [18].

To study reaction diversity across the EC classification, we developed a method based on the 42 multi-reaction isomerase EC numbers to automatically label the type of diversity in any multireaction EC number *(different* reactants, *generic* reaction on the basis of R-group and stereochemistry, *partial* reaction and *different* types of reactions). The strategy comprised a set of conditional statements combining bond change results from EC-BLAST, which allowed the detection of *different* types of reaction; comparisons of substrate and product structures and identification of R-groups and stereochemistry using Open Babel [37] and in-house scripts, which helped to find *generic* and *partial* reactions (S1 Fig.). Finally, manual analysis of 10% of the remaining multi-reactions EC numbers, which were not detected by the conditions addressing the other diversity groups, revealed them as cases of *different* reactants. This test reduced the bias caused by starting from multi-reaction isomerase EC numbers in the first place.

We tested the performance of the method by assessing its ability to correctly identify the type of diversity in fifty randomly-selected multi-reaction EC numbers from the whole of the EC classification. The test dataset comprised 22 oxidoreductases (EC 1), 19 transferases (EC 2), 5 hydrolases (EC 3), 2 lyases (EC 4) and 2 ligases (EC 6), which were manually assigned to a reaction diversity group allowing performance to be evaluated (S2 Fig.). The selection of test multireaction EC numbers was carried out randomly, but it was assured that it covers the whole diversity space of the EC classification. Overall, the method successfully assigned the correct diversity group in 41 of the total of 50 test EC numbers. Nine remaining cases could not be correctly assigned due to data errors, detection problems and atom-atom mapping accuracy (S1 Text).

## Results

### Relationship between EC number and reaction in isomerases

In general, the intrinsic diversity in isomerase multi-reaction EC numbers was interpreted in terms of the chemical variability between the reactions linked to the same EC number. In the context of catalytic promiscuity, previous studies defined reactions to be *different* if they differ in the types of bond changes (formed and cleaved), the reaction mechanism or both [38,39]. The reactions associated with the 42 multi-reaction isomerase EC numbers were manually analysed on the basis of bond and stereochemistry changes and EC numbers were divided into three groups according to *same, partial* and *different overall* chemistry of the reaction (Fig. 2). According to our observations, the first group was then further divided into two subgroups: *different* reactants and *generic* reaction. Since the EC number only describes the *overall* reaction, we do not include mechanisms in this analysis. Below is an explanation of each subgroup.

**Fig. 2.**
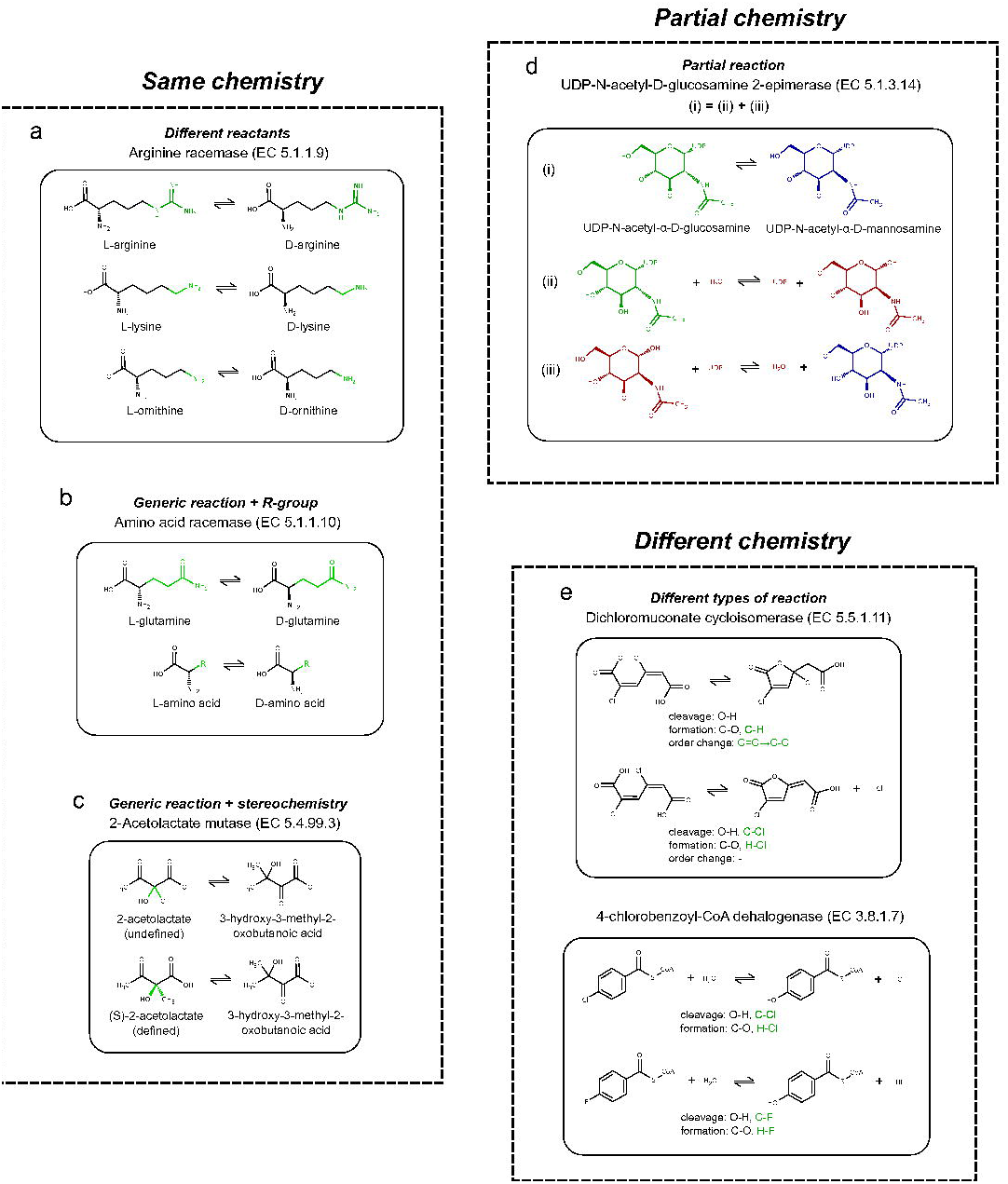
Examples of isomerase EC numbers associated with more than one enzyme reaction. (a) Arginine racemase (EC 5.1.1.9) is an isomerase acting on *different* reactants. The variability in chemical substituents is highlighted in green and the common scaffold in black. (b) Amino acid racemase (EC 5.1.1.10) is an example of *generic* reaction on the basis of R-group. Same colouring as in (a). (c) 2-acetolactate mutase (EC 5.4.99.3) is an example of *generic* reaction based on stereochemistry. The stereochemistry of C2 in acetolactate is represented as straight (undefined), up and down (defined) bonds and highlighted in green. (d) UDP-N-acetyl-D-glucosamine 2-epimerase (EC 5.1.3.14) belongs to *partial* reaction, (i) *overall* reaction – epimerisation of UDP-N-acetyl-α-D-glucosamine (green) and UDP-N-acetyl-α-D-mannosamine (blue), (ii) first *partial* reaction – hydrolysis and epimerisation of UDP-N-acetyl-α-D-glucosamine and (iii) second *partial* reaction – addition of UDP to N-acetyl-α-D-mannosamine. Intermediate compounds are highlighted in red. (e) Dichloromuconate cycloisomerase (EC 5.5.1.11) and 4-chlorobenzoyl-CoA dehalogenase (EC 3.8.1.7) catalyse *different* types of reactions. Shared bond changes are coloured in black, whereas different bond changes in green.

In the *different* reactants subgroup, reaction diversity arises due to the presence of different chemical substituents on a common structural scaffold. For example, the so-called “arginine racemase” (EC 5.1.1.9) describes the racemisation of arginine, lysine and ornithine. The three reactions involve a chiral inversion of the common Ca in the amino acid (Fig. 2a).

*Generic* reactions are used to represent multiple reactions by means of the chemical composition of their reactants. They are represented using Markush labels (e.g. R-groups) [40], which serve as chemical wildcards for other reactions. Almost one in five EC numbers are associated to at least one *generic* reaction, half of them refer to multi-reaction EC numbers and the other half represent single-reaction EC numbers (Fig. 1a). Although the association between Markush labels from the *generic* reaction and the corresponding chemical substructures in exemplar reactions is direct for multi-reaction EC numbers, this correspondence in single-reaction EC numbers is challenging where comparisons with all the other EC numbers are required.

Multi-reaction EC numbers where at least one reaction is *generic* are the subject of this study. We found that *generic* relationships according to chemical composition are of two types. First, some cases resemble the characteristics of the *different* reactants subgroup but the various chemical substituents are collectively displayed in an additional *generic* reaction, which represents the rest of reactions. For instance, amino acid racemase (EC 5.1.1.10) is linked to five reactions. Four of them describe racemisations of glutamine, serine, ornithine and cysteine and the extra one represents all of them by encapsulating the diversity of the amino acid side chain into a R-group (Fig. 2b). In some cases however, the *generic* reaction is the common structural scaffold shared among all reactions. As a result, there is no R-group involved, and the reactants of the *generic* reaction are substructures of the reactants of the rest of reactions. For example, in Fig. 2a the reactants in the epimerisation of L-ornithine are substructures of the reactants in the epimerisation of L-arginine, hence the former could also be a *generic* reaction of the latter. Although the latter *generic* relationship is evident in our manual analysis, in the process of developing an automatic method to assign EC numbers to reaction diversity groups (see Automatic analysis section) we considered this as an example of *different* reactants. Other isomerase EC numbers fall into this category such as chalcone isomerase (EC 5.5.1.6), which catalyses reversible cyclisation of chalcone into flavanone as common structural scaffold. In addition, it also performs the same reaction in hydroxy-substituted derivatives of chalcone and flavanone [41].

The second case of representation by *generic* reaction arises due to differences in the definition of stereochemistry between the *generic* reaction and rest of the reactions. Here, undefined stereochemistry (in the form of wiggly or non-stereo bond) characterises one of the chiral carbons in the *generic* reaction, whereas stereochemistry is defined for that atom in the rest of the reactions. Although a previous study reported data challenges due to the lack of stereochemical completeness in KEGG metabolites and reactions [42], to some extent recent versions of the database have incorporated these recommendations to improve the handling of stereochemistry and related data inconsistencies. Taken together, the common existence of cases of defined and undefined stereochemistry in several EC numbers supported the formulation of this diversity group. For example, acetolactate mutase (EC 5.4.99.3) is associated with two reactions: the isomerisations of 2-acetolactate (generic reaction, undefined stereochemistry) and (S)-2-acetolactate (specific reaction, defined stereochemistry) (Fig. 2c). As in *generic* reactions on the basis of R-group, cases of undefined stereochemistry in the form of wiggly bonds were detected in our automatic method, however the cases of non-stereo bonds were regarded as examples of *different* reactants.

It is a well known fact that there are enzymes releasing intermediate products of an *overall* reaction from the active site [5]. Reactions leading to these intermediates are known as *partial* reactions. Similarly, an enzyme may subsequently catalyse two or more *partial* reactions with or without releasing any intermediates, these are considered as *consecutive* reactions. For example, in Fig. 2d UDP-N-acetyl-D-glucosamine 2-epimerase (EC 5.1.3.14) catalyses the epimerisation of UDP-N-acetyl-α-D-glucosamine and UDP-N-acetyl-α-D-mannosamine (*overall* reaction). This transformation comprises two successive *partial* reactions in the mechanism – hence, they are *consecutive.* First, the UDP moiety is hydrolytically eliminated from the anomeric carbon and epimerisation takes place at C2 (first *partial* reaction). Second, the UDP moiety is added to the anomeric carbon (second *partial* reaction). Combining these two *consecutive* reactions leads to the *overall* reaction. Whereas this example summarises this group in its simplest form, we also found three other alternatives of *partial* reactions linked to the same EC number, which are described in S1 Text. Previous studies have alternatively used the concept of “multi-step reaction” to refer to our definition of *overall* reaction composed of more than one *partial* reactions that occur consecutively [6]. However, the term step in a reaction usually implies one mechanistic step of the *overall* reaction. As mechanisms are not included in the EC classification, we preferred using the term *partial* reaction in order to avoid confusion.

Finally, EC numbers might also be linked to at least two *different* types of reactions. Dichloromuconate cycloisomerase (EC 5.5.1.11) catalyses two types: first, the isomerisation of 2,4-dichloro-cis,cis-muconate and 2,4-dichloro-2,5-dihydro-5-oxofuran-2-acetate and also, the conversion of 2,4-dichloro-cis,cis-muconate into trans-2-chlorodienelactone and chloride (Fig. 2e) [43,44]. Although the two reactions share the cleavage of O-H and formation of C-O bonds, they differ in other bond changes, so they are considered to be *different.* However the product of the first isomerisation might eliminate chloride to yield trans-2-chlorodienelactone in an uncatalysed manner and therefore the second reaction would be the result of an isomerisation and successive elimination, which can also be interpreted as an example of *partial* reaction as described before. Other examples of EC numbers that can also be categorised under both *different* types of reaction and *partial* reaction involve sugar isomerisations such as those catalysed by D-arabinose isomerase (EC 5.3.1.3) and ribose-5-phosphate isomerase (EC 5.3.1.6) where the ring opening and closure might be uncatalysed. Perhaps a more definite example of *different* reaction types is 4-chlorobenzoyl-CoA dehalogenase (EC 3.8.1.7). This EC number involves the dehalogenation of 4-chlorobenzoyl-CoA into 4-hydroxybenzoyl-CoA and also the hydrolysis of the fluoro, bromo and iodo derivatives (Fig. 2e). This can also be interpreted as an example of *different* reactants with a halogen atom corresponding to a *generic* substructure.

Following our manual classification, 30 of the 42 multi-reaction isomerase EC numbers were solely assigned to one of the groups, whereas the diversity of the remaining 12 EC numbers was explained by more than one group. Overall, 57 group assignments were manually designated: 24 *different* reactants, 17 *generic* reactions (R-group and stereochemistry), 5 *partial* reactions and 11 *different* types of reactions. Among the EC numbers assigned to more than one group, we found 2-acetolactate mutase (EC 5.4.99.3) (Fig. 2c). In addition to the transfer of a methyl group from C2 to C3 in (S)-2-acetolactate, this isomerase also catalyses the transfer of an ethyl group from C2 to C3 in (S)-2-aceto-2-hydroxybutanoate. This EC number could be assigned to both groups: *generic* reaction on the basis of stereochemistry and *different* reactants (S3 Fig.). Similarly, although dichloromuconate cycloisomerase (EC 5.5.1.11) is an example of *different* types of reactions (Fig. 2e), a potentially uncatalysed elimination of chloride may also link these two reactions in a *partial* relationship.

### Relationship between EC number and reaction in the EC classification

A schematic diagram illustrating the various groups of reaction diversity is shown in Fig. 3a. There are 1277 multi-reaction EC numbers in the entire EC classification, 90% of them (1153) could be analysed using our method. The most common group was *different* reactants including almost half of the examples. *Different* reaction types followed with 29% and ultimately *partial* and *generic* reactions made up the rest (Fig. 3b). The overall distribution was similar in oxidoreductases (EC 1), transferases (EC 2) and hydrolases (EC 3), which were correspondingly the EC classes involving the highest number of multi-reaction EC numbers (Fig. 3c) and not surprisingly, also the EC classes with the largest number of EC numbers in the EC classification [45]. Exceptionally, the most common diversity group in ligases (EC 6) is *different* reaction types, instead of *different* reactants. Also, the method did not identify any example of EC numbers involving *generic* reactions in lyases (EC 4) and ligases (EC 6).

**Fig. 3.**
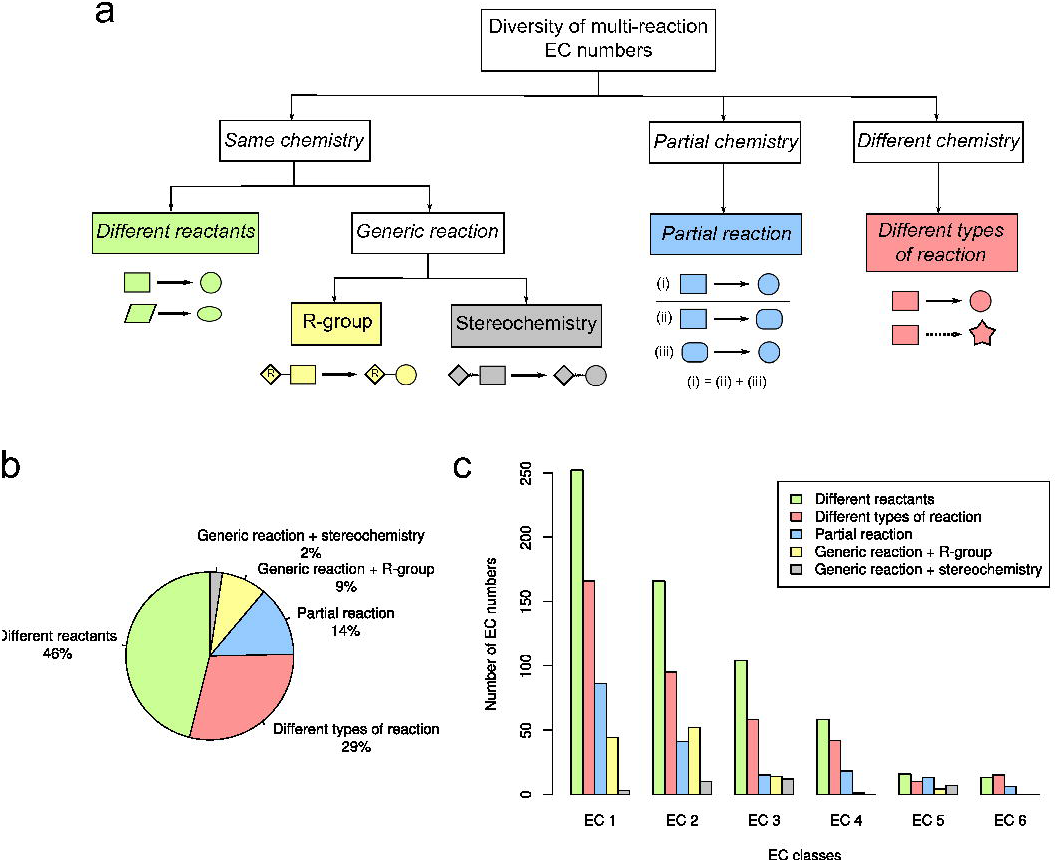
An overview of reaction diversity in the EC classification. (a) A schematic diagram summarising the groups of reaction diversity. (b) Frequency of reaction diversity group assignments. (c) Total number of multi-reaction EC numbers by EC class for each group of reaction diversity.

## Discussion

### Overall

Although there is literature reported by the IUBMB discussing specific cases of reaction diversity across the EC classification [5], the aim of this study was to systematically explore aspects of the chemical diversity in the description of enzyme function in a specific EC primary class manually and automatically for the entire EC classification. In order to extract bond changes from reactions we used the EC-BLAST algorithm, which is based on chemical concepts, such as the principle of minimum chemical distance and chemical bond energies, in order to guide the atom-atom mapping and chemical matrices for similarity searches [18]. As suggested in a recent review [46], the incorporation of chemical knowledge adds accuracy to existing strategies to perform reaction comparison.

This study depends on the quality of reaction data available in the KEGG database [42]. We found this to be the major source of discrepancy between the manual and automatic analyses since many reactions were not balanced hence consistent atom-atom mapping becomes impossible. Whereas multiple strategies to correct unbalanced reactions [46-48] and to reconcile biochemical reactions across databases [34] have been recently presented, novel improvements of the algorithms and further data curation and integration are needed [49,50]. In addition, the quality of the manual curation performed in this study is dependent on the authors’ ability to interpret reactions, as well as the experimental information available in the primary literature. The automatic analysis relied only upon the overall reaction equation and the ability of EC-BLAST to compute accurate atom-atom mappings.

To what extent do the findings of this study overlap with those discovered in previous accounts on enzyme promiscuity? There are obviously enzymes catalysing different reactions with different EC numbers, but the IUBMB does not usually include this for most enzymes. However, to some degree, the working definitions of substrate and product promiscuity [51] somewhat resemble our diversity groups of *different* reactants and *generic* reactions. Likewise, catalytic promiscuity partly corresponds to *different* reaction types. However, whereas promiscuity definitions are genuinely attributed to enzymes in order to describe their ability to catalyse more than one reaction, our characterisation of reaction diversity applies to diversity within the same EC number, which adds an extra level of chemical variability to the existing definitions of enzyme function.

The surprising observation of this study is that almost one-third of the EC numbers involving more than one reaction have *different* reaction types, bearing key differences in catalysed bond changes. Whereas some of them also correspond to *partial* reactions, many are cases of catalytic promiscuity within the same EC number where the annotated enzyme catalyses two or more distinct reactions. Manual analysis revealed that most cases are similar to 4-chlorobenzoyl-CoA dehalogenase (EC 3.8.1.7) (Fig. 2e) indicating that whereas some bond changes are shared, the rest individually characterise each of the different reactions.

The rationale behind why the IUBMB and reaction databases have assigned multiple biochemical reactions to the same EC number is to some extent comprehensible. For instance, the product of some catalysed reactions sometimes undergoes a fast and uncatalysed reaction while still in the active site. These EC numbers comprise two reactions: one comprising only the catalysed reaction and another consisting of the catalysed+uncatalysed *consecutive* reactions. Whereas some enzymologists might preferably associate the EC number only with the catalysed reaction, the fact that the uncatalysed reaction takes place in the enzyme’s confinement supports the catalysed+uncatalysed interpretation (see Experimental and Results).

However the complexity in the relationship between reaction and EC number goes beyond this study and cases of *generic* relationships are also common in single-reaction EC numbers (Fig. 1a) and across different EC numbers. For example, as highlighted before, EC 5.1.1.10 was defined by the IUBMB after the discovery of an enzyme that broadly catalyses racemisations of several amino acids [52]. The biochemical reaction contains an R-group and it effectively represents reactions catalysed by specific amino acid racemases, which are also assigned different EC numbers, e.g. alanine (EC 5.1.1.1) and serine (EC 5.1.1.18). Although this and other examples [33] were attempts to incorporate an enzyme property such as substrate specificity to guide the EC classification, this might lead in some cases to EC numbers being embedded into one another and no longer chemically independent from each other, which adds further complications to a classification based solely on the chemistry of the overall reaction.

### Improving the description of complex enzyme reactions

The ability of the IUBMB to manually update the EC classification in the form of transferred and deleted entries when new enzyme data becomes available is necessary. For example, during the fifty years succeeding the creation of the EC entry for phosphoglycerate mutase in 1961 (EC 5.4.2.1), evidence supporting two distinct mechanisms concerning different usage of the cofactor 2,3-diphosphoglycerate by this enzyme accumulated in the literature [53]. In 2013, the original EC number was transferred to EC 5.4.2.11 (cofactor-dependent) and EC 5.4.2.12 (cofactor-independent). In addition, several expert recommendations concerning definition and handling of EC numbers in biological databases have already been suggested in different contexts. For example, Green and Karp advised about the problems associated with the assignment of partial EC numbers (those containing a dash, e.g. EC 5.1.1.-) to genes and proposed changes to the specification of these ambiguous identifiers [54]. Similarly, we suggest approaches to clarify multi-reaction EC numbers, which will hopefully help to improve the EC and reaction databases [5] and serve to guide standards for the reporting of enzyme data [55-57] and existing initiatives for the assignment of enzyme function [58-60].

A multi-reaction EC number belonging to the groups’ *different* reactants or *generic* reactions could either be combined into a single-reaction EC number (<u>collective</u> approach) or split into as many distinct EC numbers (<u>specific</u> approach). In the first place, diversity could be represented by R-group definitions, which would encapsulate chemical substituents at different positions in the reactants. When necessary, stereochemically-undefined bonds could also be employed to indicate the non-stereoselectivity of some biochemical reactions (Fig. 4a). Secondly, the <u>specific</u> strategy arises when there are significant changes of substrate specificity between enzymes annotated with the same multi-reaction EC number. Instead of defining a *generic* reaction, it might be more sensible to re-define several EC numbers according to the distinct patterns of substrate specificity [61]. However, although EC-BLAST provides a robust method to measure chemical differences between overall reactions in a continuous manner, defining the cut-offs required to designate separate EC numbers (for example, between different substrates) is *a priori* arbitrary and would need to be addressed explicitly.

**Fig. 4.**
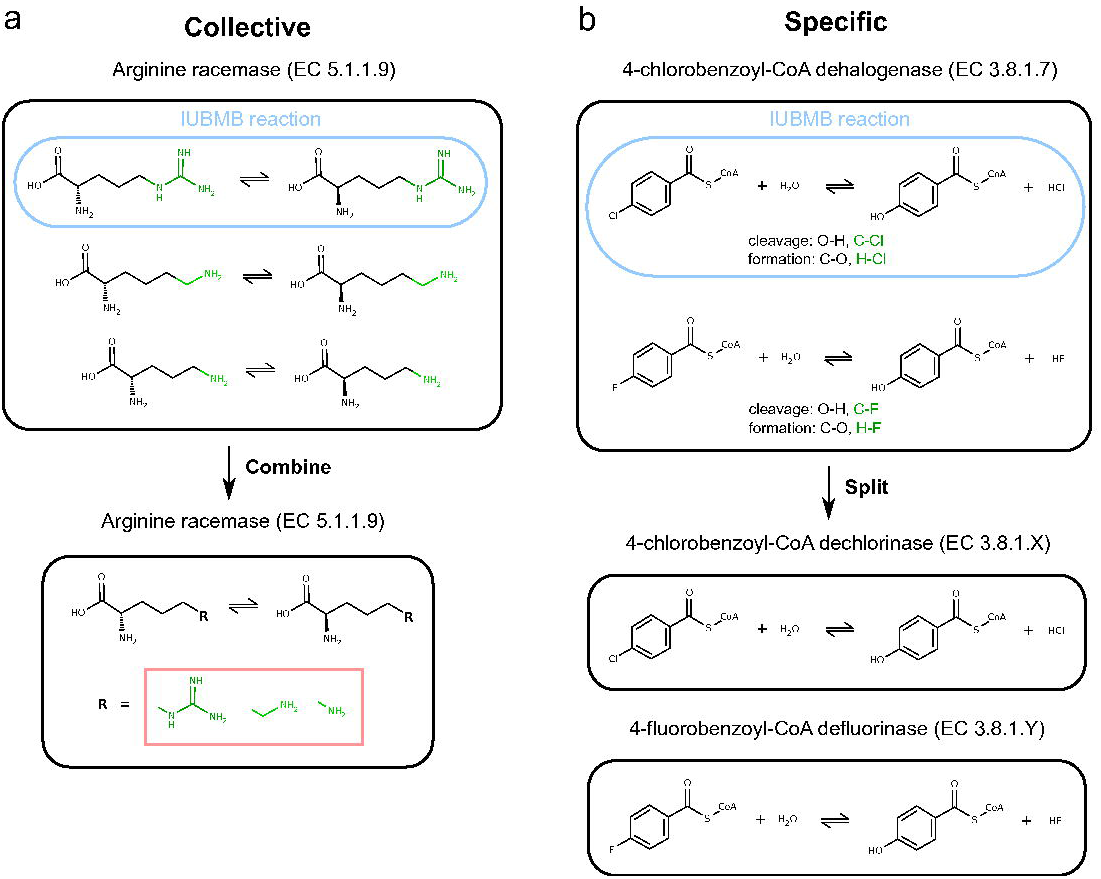
Examples of the <u>collective</u> and <u>specific</u> approaches. (a) The *different* reactants of arginine racemase (EC 5.1.1.9) are combined into a single-reaction EC number using R-group. (b) The two *different* types of reaction catalysed by 4-chlorobenzoyl-CoA dehalogenase (EC 3.8.1.7) are split and re-defined into two single-reaction EC numbers.

A proposed *modus operandi* when dealing with *different* reaction types involves using the <u>specific</u> approach to divide the multi-reaction EC number into multiple EC numbers, one for each *different* reaction [27] (Fig. 4b). Regarding *partial* reactions, we recommend to collectively reduce the multireaction EC number by combining all *partial* reactions with required enzyme catalysis into a singlereaction EC number, while setting uncatalysed reactions aside.

Both <u>collective</u> and <u>specific</u> approaches have several benefits. For instance, three main advantages characterise the <u>collective</u> approach. First, it is a compact way to arrange reaction information in a clear and structured manner. Second, it conveys how chemists and biochemists represent reactions in the literature, databases and patents [62-64]. Third, diversity can be captured using Markush labels such as R-groups [40,65], which would be subsequently described in associated files, tables or chemical libraries [66]. Alternatively, diversity in the reactants could be encoded using recent developments in the description of chemical patterns [67]. Also, the <u>collective</u> approach brings together reactions that are often evolutionarily-related. The precise definition of R-groups will also help previous studies that were limited in their ability to handle *generic* structures. Although some strategies did not explicitly define R-groups in their representation of biochemical reactions [68], several studies preprocessed oxidoreductase (EC 1) and hydrolase (EC 3) reactions by replacing every R-group by a hydrogen atom [8,69] or methyl group [70] in order to calculate physicochemical and topological properties in atoms and bonds involved in reaction centres. Using more specific substitutions, R-groups were manually replaced by methyl, adenine, cytosine or other chemical moieties depending on the type of biochemical reaction [30,31]. These studies suggest that having EC number-specific definitions of R-groups based on experimental evidence is a necessary step in order to implement the <u>collective</u> approach across the classification.

Whereas the <u>collective</u> approach relies on presenting a common structural scaffold and diversity encoded as chemical placeholders, the <u>specific</u> approach is divisive and explicitly distinguishes between reactions that are considered as chemically distinct. A clear advantage of the latter is when subtle differences between biochemical reactions are captured using different EC numbers, for instance, distinct bond changes or substrate specificity. The description of enzyme function will then be more detailed and it will help to dissect some of the complexities in the relationship between enzyme sequence, structure and function [10].

The terms of the application of the <u>collective</u> and <u>specific</u> approaches to combine or split multireaction EC numbers are proposed in the following recommendations to improve the description of multi-reaction EC numbers:

- Reactions sharing the *same overall* chemistry (identical bond changes) should be combined into a single-reaction EC number (corresponding to groups: *different* reactants and *generic* reaction). The chemical diversity observed as different embodiments of a *generic* structure would be encapsulated using R-group definitions and stereochemically-undefined bonds in associated libraries and chemical patterns.
- If reactions have *different overall* chemistry (distinct bond changes), the EC number should be split in multiple single-reaction EC numbers (group: *different* types of reaction). Similarly, reactions catalysed by enzymes annotated with the same EC number that display distinct substrate specificities or cofactor dependencies should also be split in as many single-reaction EC numbers as patterns of specificity exist (groups: *different* reactants and *generic* reaction).
- Reactions sharing *partial overall* chemistry (several *partial* reactions integrate into an *overall* reaction) should be treated carefully. The *partial* reactions that take place in the active site of the enzyme should be combined into a single-reaction EC number (group: *partial* reaction) with chemical diversity encapsulated in libraries as described before. Uncatalysed *partial* reactions should be considered separately.

As a way to summarise the diversity existing in a multi-reaction EC number, biological databases such as KEGG [2] rely on the so-called “IUBMB reaction”. This is the reaction assigned to the EC number by the IUBMB in the first place, which is chosen by KEGG as the representative reaction for the group of reactions associated with the same EC number (Fig. 4). Whereas this assignment is useful when selecting an example reaction from an EC number and it was adopted as a principle in the development of other reaction databases such as Rhea [71], it is sometimes missing or conflicting and it also overlooks the existing diversity. For instance, EC 5.1.1.13 is described as “Reaction: L-aspartate = D-aspartate” and “Comments: Also acts, at half the rate, on L-alanine”, which is a rather vague description. Similarly, some EC numbers are not associated to any IUBMB reaction and also, EC numbers are sometimes linked to the same IUBMB reaction, 2,3-diphosphoglycerate-dependent and independent phosphoglycerate mutases (EC 5.4.2.11 and EC 5.4.2.12) are both assigned the same IUBMB reaction comprising the isomerisation of 2-phospho-D-glycerate to 3-phospho-D-glycerate. Taken together, from the authors’ perspective, a more robust and consistent approach to describe multi-reaction EC numbers is needed.

This systematic analysis is relevant for the functional annotation of sequenced genomes and by extension, it has implications for our ability to build and compare genome-scale metabolic reconstructions [72-74]. There is a direct correspondence between EC numbers and terms representing the molecular function of protein-coding genes in the Gene Ontology (GO) [75], which implicitly adopted EC numbers as part of their classification. This ontology is currently the widely used standard for the automatic assignment of function to proteins and genes [76]. We observed that multi-reaction EC numbers/GO terms are commonly transferred between similar enzymes during this process. Such a predicted assignment of function does not consider that enzymes annotated with the same multi-reaction EC number might have different reaction specificities in different species, which may lead to a general overestimation of the catalytic capabilities of organisms as predicted from their genomes.

## Conclusions

To summarise, this study adds an additional level of chemical complexity to our current description of enzyme function using EC numbers. Remarkably, almost a third of all known EC numbers are associated with more than one enzyme reaction in the KEGG database. Existing approaches to handle this diversity are ineffective, therefore we decomposed this diversity into four categories: *different* reactants, *generic*, *partial* and *different* types of reaction with the aid of computational methods to automatically compare reactions. All multi-reaction EC numbers in our database, annotated according to our reaction typing are given in S1 Table. We hope this information will help to improve our understanding and description of enzyme reactions.

## Acknowledgements

Authors acknowledge KEGG for making their reaction data available for academic use through their API services. SMC acknowledges Dr. John BO Mitchell for critically reading the manuscript.

## Supporting Information

**S1 Text. Extension of the methods and results described in the manuscript.**

**S1 Fig. Workflow illustrating the automatic analysis of multi-reaction EC numbers.**

**S2 Fig. Results of the test to evaluate the automatic method labelling multi-reaction EC numbers according to the reaction diversity group.**

**S3 Fig. 2-Acetolactate mutase (EC 5.4.99.3) is an example of EC number assigned to two groups of reaction diversity: *different* types of reaction and *partial* reactions.**

**S1 Table. Table listing all the multi-reaction EC numbers considered in this study.** They have been annotated according to our description of chemical diversity groups and isomerase EC numbers have been manually labelled with our recommendation for improvement.

